# Predicting the evolution of adaptation and plasticity from temporal environmental change

**DOI:** 10.1101/2023.02.12.528221

**Authors:** Cristóbal Gallegos, Kathryn A. Hodgins, Keyne Monro

## Abstract

Environmental change drives evolutionary adaptation, which determines geographic patterns of biodiversity. At a time of rapid environmental change, however, our ability to predict its evolutionary impacts is far from complete. Temporal environmental change, in particular, often involves joint changes in major components such as mean, trend, cyclic change, and noise. While theoretical predictions exist for adaptation to temporal change in isolated components, knowledge gaps remain. To identify those gaps, we review the relevant theoretical literature, finding that studies rarely assess the relative effects of components changing simultaneously, or attempt to translate theoretical predictions to field conditions. To address those gaps, we draw on classic evolutionary theory to develop a model for the evolution of environmental tolerance, determined by an evolving phenotypically plastic trait, in response to major components of temporal environmental change. We assess the effects of different components on the evolution of tolerance, including rates of adaptation towards new environmental optima, and the evolution of plasticity. We retrieve and synthesize earlier predictions of responses to components changing in isolation, while also generating new predictions of responses to components changing simultaneously. Notably, we show how different forms of environmental predictability emerging from the interplay of cyclic change, stochastic change (noise), and generation time shape predicted outcomes. We then parameterise our model using temperature time series from global marine hotspot in southern Australia, illustrating its utility for predicting testable geographic patterns in evolved thermal tolerance. Our framework provides new insights into the evolution of adaptation and plasticity under temporal environmental change, while offering a path to improving predictions of biological responses to climate change.

## Introduction

Environmental change is ubiquitous and can shape diversity at different levels of biological organization (Ruokolainen et al., 2009; Scheffers et al., 2016). Changes in environmental temperature, for example, affect biochemical reactions within cells, organismal physiology, performance, and fitness, and species’ geographic distributions from local to global scales (Clarke, 2017; Pecl et al., 2017; Peters et al., 2016). Environmental change can also shape evolutionary processes — including selection, adaptation, speciation, and extinction (e.g., Bell & Gonzalez, 2009; Hoekstra et al., 2001; Hoffmann & Sgrò, 2011; Rescan et al., 2020; Schluter, 2001) — that should underpin biodiversity and determine its geographic distribution (Connallon & Sgrò, 2018; Polechová, 2018; Polechová & Barton, 2015). Yet, at a time of rapid and increasing climate change (Smith et al., 2015), our ability to predict its evolutionary impacts on biodiversity is far from complete (Catullo et al., 2019).

Two key modelling approaches are widely used to predict geographic patterns in biodiversity. First, niche (or species distribution) models predict species’ ranges based on correlations between occurrence and climatic conditions (Cheung et al., 2009; Elith & Leathwick, 2009; Fleishman et al., 2001), but are criticised for ignoring biological processes that dictate occurrence (Davis et al., 1998; Dormann, 2007; Hijmans & Graham, 2006; Pearson & Dawson, 2003). Mechanistic niche models address such criticism to a large extent (Kearney et al., 2009; Kearney & Porter, 2009), but remain out of reach for all but a few well-studied species (Dormann et al., 2012; Evans et al., 2015). Moreover, recent efforts to incorporate adaptation (e.g., Bush et al., 2016; Catullo et al., 2019; Diamond, 2018; Martínez-Padilla et al., 2017) are hampered by limited understanding of how it varies in response to climate. Second, mechanistic population models combine evolutionary genetics and demography to predict how adaptive changes in key traits that affect population growth allow populations to persist in the face of environmental change (Burger & Lynch, 1995; Chevin et al., 2010; Lande & Shannon, 1996; Lynch & Lande, 1993; Willi & Hoffmann, 2009). Such models cannot, however, accurately predict adaptation without including environmental variables (Chevin et al., 2010) and, by extension, the effects of those variables changing at different temporal or spatial scales that can produce different evolutionary outcomes. There is thus a need to develop tools to better predict how those outcomes vary in time and space, and to explicitly link predictions to relevant patterns of environmental change in ways that are broadly applicable to different species.

Environmental tolerance curves, relating fitness to environmental variables either directly or mediated by underlying traits (Figure 1), are powerful tools for linking evolutionary outcomes to environmental change (Carscadden et al., 2020; Chevin et al., 2010; Kingsolver & Buckley, 2017; Lynch & Gabriel, 1987; Sexton et al., 2017). Combining them with climate data, for example, predicts global shifts in geographic distributions, population growth rates, and extinction risks of marine and terrestrial ectotherms in response to projected changes in temperature (Deutsch et al., 2008; Pinsky et al., 2019; Vasseur et al., 2014; Woods et al., 2018). Such insights, however, generally rest on the assumption that environmental tolerance is static in time and space, when it may instead be expected to evolve over time as populations adapt to environmental change, and to vary geographically among lineages exposed to different forms and magnitudes of change (Gabriel & Lynch, 1992; Lande, 2014; Lynch & Gabriel, 1987; Sheth & Angert, 2014; Wieczynski et al., 2018). Progress on this front is limited by our poor understanding of, and ability to predict, how complex, real-world patterns of environmental change shape environmental tolerance in nature. Given the daunting task of addressing this gap empirically, a unified theoretical framework for predicting the evolution of tolerance in the face of real-world change, and the further implications for geographic patterns in biodiversity, is therefore a pressing endeavour for evolutionary biology with clear relevance for conservation and management.

**Figure 1.**
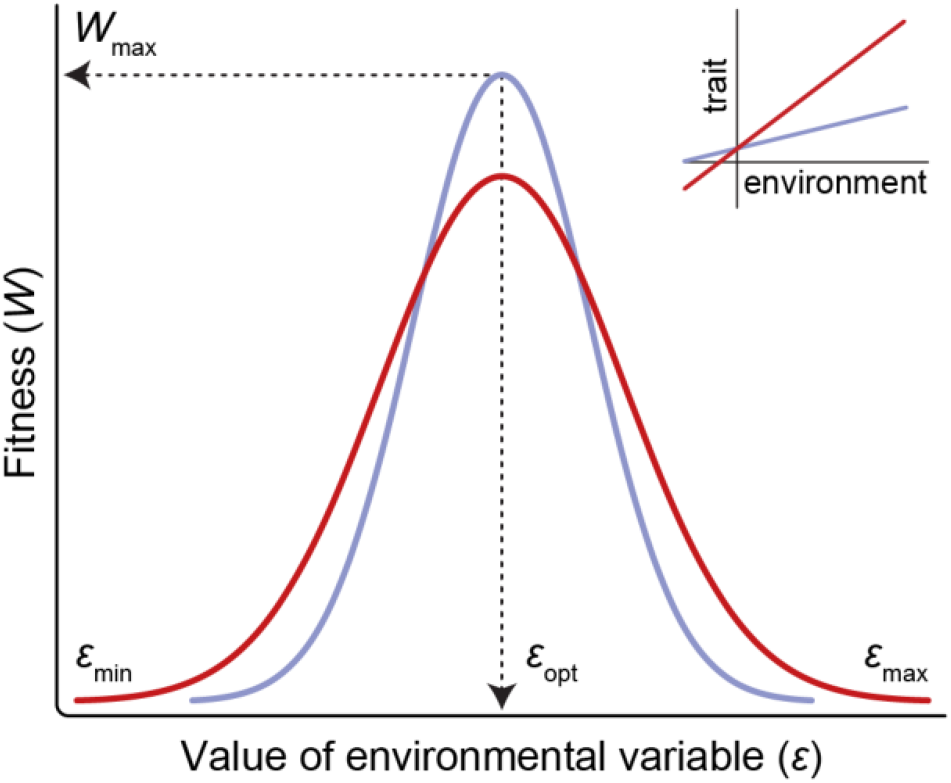
Environmental tolerance curves relating fitness (*W*) to an environmental variable (*ε*). Each curve is described by a few key terms: *ε*_*opt*_ is the environmental optimum at which fitness is maximised (*W*_max_) and *ε*_*min*_ and *ε_max_* are the lower and upper environmental limits at which fitness declines to zero. Tolerance breadth is then the range of environmental values over which fitness is positive, comparable to the fundamental niche. Breadth is determined by phenotypic plasticity in an underlying trait, described by the slope of the linear reaction norm relating trait expression to the environment (inset). The red tolerance curve is broader than the blue tolerance curve due to greater plasticity in the trait, but also has a lower peak due to the cost of plasticity.

Temporal change in any environmental variable can be decomposed into different components — namely, an abrupt change in the mean, constant directional change (or trend) in the mean, cyclic change in values around the mean, and stochastic change (or noise) unexplained by other components (Figure 2; Chatfield, 1996; Ruokolainen et al., 2009; Vasseur & Yodzis, 2004). These components are expected to have different impacts on the evolution of environmental tolerance. To the extent that a variable affects fitness, its trend imposes directional selection through sustained, directional change in the optimum phenotype. In the absence of constraints (e.g., lack of suitable genetic variation), a population’s mean phenotype should evolve to track the moving optimum (Lande & Shannon, 1996), thereby shifting the peak of the environmental tolerance curve (*ε*_*opt*_ in Figure 1) towards lower or higher tolerance (left or right on the environmental axis in Figure 1). Trend remains by far the best-studied component of environmental change due to the pivotal role of directional selection in adaptation and diversification (Kingsolver & Diamond, 2011; Rieseberg et al., 2002), but calls are mounting to integrate other components into predictions (Bates et al., 2018; Dillon et al., 2016; Dowd et al., 2015; Helmuth et al., 2014). In theory, environmental cycles with greater amplitudes may broaden environmental tolerance by promoting the evolution of adaptive phenotypic plasticity (Figure 1; Gilchrist, 1995; Ketola et al., 2013; Lynch & Gabriel, 1987; Scheiner, 1993), whereas greater environmental noise may inhibit the evolution of adaptive tracking or adaptive plasticity (Lande & Shannon, 1996; Reed et al., 2010). Populations in highly-stochastic environments may thus be prone to genetic drift, risking loss of genetic diversity and fitness (Lande & Shannon, 1996; Tufto, 2015). In general, however, we still lack a thorough understanding of how different components of environmental change combine to shape adaptation and plasticity, and their rates of evolution, across populations and landscapes (Catullo et al., 2019; Rescan et al, 2022; Shaw & Etterson, 2012; Vinton et al., 2022; Visser, 2008). This understanding is crucial to predicting adaptive capacity and persistence under the fast-paced and complex nature of climate change.

**Figure 2.**
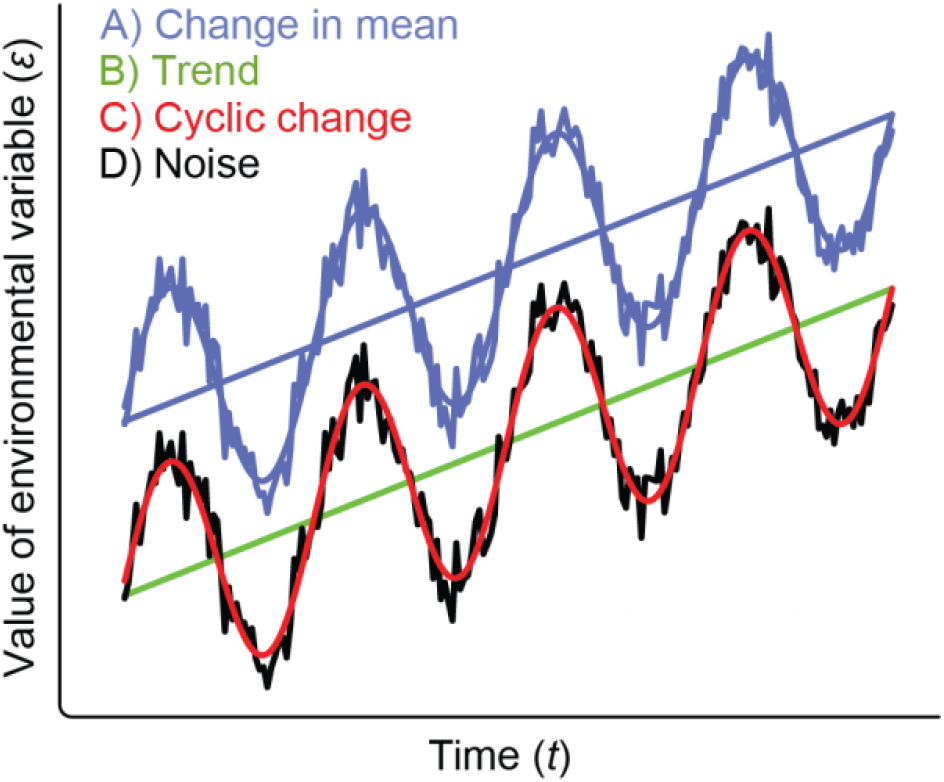
Temporal change in an environmental variable (*ε*) can be decomposed into different components. A) Change in mean is an abrupt shift in the mean environmental value (or intercept) at a given time (*t*), which organisms can face *in situ* or as they disperse. B) Trend is a constant rate of directional change in the mean environmental value, described by the slope of the line relating mean values to time. C) Cyclic change is predictable variation in environmental values around the mean, as in daily or seasonal variation in temperature. Cycles are described by sine waves of given amplitude (deviation from the mean) and period (time between repeated peaks or troughs), or by summary statistics such as range. D) Noise is stochastic change left in environmental values after subtracting other components, and can also vary in structure (known as noise colour). Noise colour is described by *ρ*, the autocorrelation of noise, or *β*, the exponent of the power law *s*(*f*) = 1/*f*^*β*^, where *f* is the frequency of noise and *s*(*f*) is its spectral density (in practice, *β* is estimated as the negative slope of the line relating log _l0_ frequency to log _l0_ spectral density). Noise in most environments ranges from white (*ρ* ≈ 0 or *β* ≈ 0), whereby values are temporally uncorrelated and hence unpredictable, to reddened (*ρ* > 0 or *β* > 0), whereby changes are positively correlated and hence more predictable. See Chatfield (1996), Ruokolainen et al. (2009), and Vasseur & Yodzis (2004) for details. Note that components are shown additively (e.g., the black line includes trend, cyclic change, and noise).

Here, we start with a brief, targeted review of the theoretical literature on adaptation to temporal environmental change (Supporting Information 1), to identify key knowledge gaps in the evolution of environmental tolerance. We find that most theoretical models do not explore how multiple components of change jointly affect adaptation, and are rarely parameterized with real environmental data in order to translate model predictions to conditions in nature (but for the latter see Chevin et al., 2015). To address these gaps, we draw on existing evolutionary theory (Chevin & Lande, 2010; Chevin et al., 2010; Lande, 1976, 2009, 2014) to develop a quantitative genetic model for the evolution of environmental tolerance, via evolution of an underlying plastic trait that determines fitness, in response to joint changes in environmental mean, trend, cycle, and noise (Supporting Information 2). We retrieve and synthesize earlier predictions of responses to isolated changes in these components, while also providing new insights into the relative effects of components acting simultaneously. Notably, we show how different forms of environmental predictability emerging from the interplay of cyclic change, noise, and generation time (relative to cycle period) shape the evolution of plasticity and tolerance. Last, we parameterise our model using real temperature time series to illustrate its utility in predicting differences in evolved thermal tolerance across a global hotspot of marine biodiversity and warming in southern Australia (Costello et al., 2022; Hobday & Pecl, 2014). Our framework combines evolutionary theory, temporal environmental change, and climate data in a way that, to our knowledge, has not been done before, and generates testable null predictions about environmental tolerance in natural populations. Better integrating evolutionary models with environmental data is a key step for advancing our understanding of population responses to climate change and predicting changes in geographic patterns of biodiversity.

### Adaptation to temporal environmental change: a brief review of theory

In parallel with accelerated climate change, interest in understanding and modelling evolutionary adaptation to temporally changing environments has grown rapidly in recent decades (Figure 3A). To identify knowledge gaps and opportunities for further developing this field, we present a targeted review of the theoretical literature. Studies were identified based on previous knowledge and additional steps (e.g., scanning citations in relevant articles; Table S1). We acknowledge this is not a systematic review of adaptation to environmental change (for which we recommend Hoffmann & Sgrò, 2011 or Kopp & Matuszewski, 2014), but rather a targeted review of theoretical modelling of adaptation to temporal environmental change specifically. Experimental studies, studies that modelled demographic but not evolutionary responses, and studies focusing on spatial environmental change, were excluded. We start by outlining the main ways in which adaptation to temporal change has been modelled. Next, we describe two interrelated themes that motivated this modelling. Last, we identify knowledge gaps that we believe could provide important insights if addressed, and improve our capacity to predict population responses to climate change.

**Figure 3.**
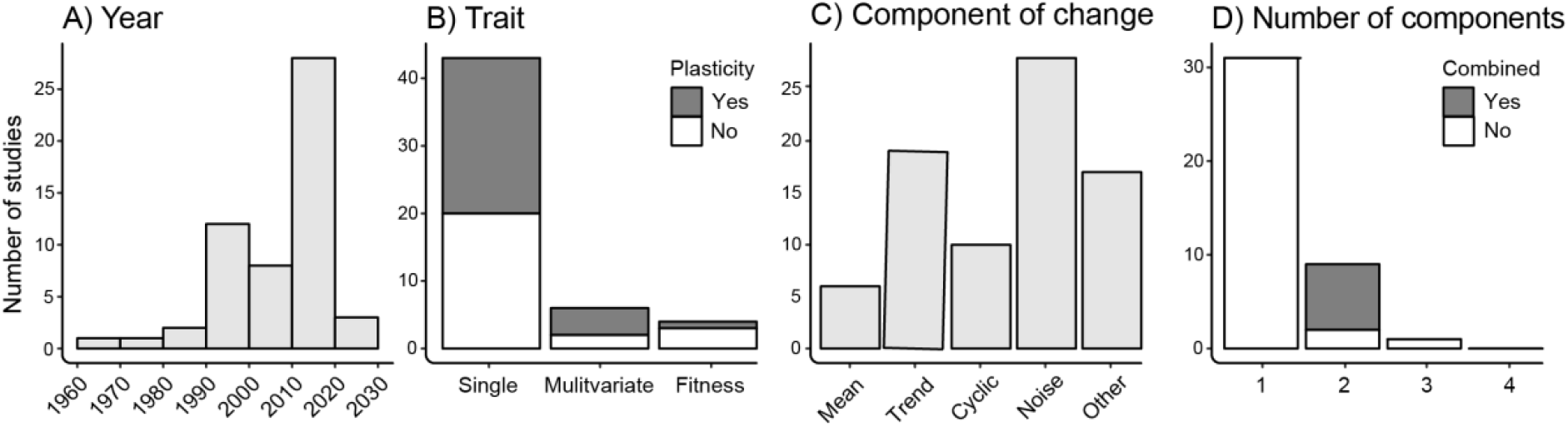
Relevant theoretical studies of adaptation to temporal environmental change in recent decades (55 studies identified from 1960s to present). A) Yearly number of studies modelling adaptation to a temporally moving optimum. B) Number of studies modelling evolution of a single trait, multivariate trait, or fitness, with plasticity (dark bars) or without plasticity (white bars). C) Number of studies modelling evolution under different components of temporal environmental change (abrupt change in mean, trend, cyclic change, noise; see Figure 2). D) Number of studies that included one, two, three, or four of those components, in combination (dark bars) or in isolation (white bars).

We identified 55 relevant studies spanning nearly six decades (Table S2), from Levins (1965) to Chevin et al. (2022). Most studies focused on selection on a single trait that determines fitness, though a few considered multiple correlated or uncorrelated traits, or evolution of fitness itself (Figure 3B). Around half of the studies included evolution of trait plasticity (Figure 3B), usually modelled as the slope of a linear reaction norm (but see Gavrilets & Scheiner, 1993), while some studies assumed plasticity to be fixed rather than evolving (e.g., Chevin et al., 2010; Hangartner et al., 2022; Marshall et al., 2016). Of studies that included plasticity, most modelled plasticity in traits determined by the environment during development and fixed for the rest of the life cycle. Some studies modelled plasticity in labile traits adjusted by the environment throughout life (as in some physiological or behavioural traits; Heatwole & Fulton, 2013; Taff & Vitousek, 2016), while a few studies modelled plasticity of both types. In terms of the nature of temporal environmental change, most studies modelled it as an abrupt change in mean, as a continuous and predictable change in mean (a constant trend or deterministic cycle), and/or as noise of varying colour (Figure 3C). However, a number of studies modelled other types of change, including multiperiodic cycles (Lande, 2019), pulsed extremes (Lyberger et al., 2021), and random or periodic alternations between states (Gabriel et al., 2005) (Figure 3C). Importantly, of the major components of temporal change in Figure 2, most studies modelled only one, some modelled two, a few modelled three, and none modelled all four (Figure 3D). Moreover, the only studies to evaluate components of temporal change in combination considered only two of them (usually trend and cyclic change, or trend and noise).

A large fraction of the models surveyed sought to understand the evolutionary causes and consequences of plasticity. This is a major theme in evolutionary ecology and reviewed amply elsewhere (e.g., Hendry, 2016; Pfennig, 2021; Schlichting, 2021), so we focus only on aspects of particular relevance here. Most studies agree that environmental variability and predictability favour the evolution of plasticity (Ezard et al., 2014; Lande, 2014; Reed et al., 2010; and references therein). However, predictability is often defined in different ways, which can greatly affect theoretical predictions (Bitter et al., 2021; Marshall & Burgess, 2015). Specifically, predictability can refer to the extent to which populations experience deterministic cycles of change as opposed to stochastic change (Ezard et al., 2014), how well conditions at one point in time predict conditions at the next (the autocorrelation of environmental noise; Tufto, 2015), or the reliability of environmental cues (the correlation between the environment that determines plasticity and the environment of selection; Tufto, 2000; Bitter et al., 2021). These different forms of predictability are interrelated, and their importance will likely depend on the type of trait studied, yet this is usually overlooked. For example, deterministic cycles and cue reliability might be most important for developmental traits, while autocorrelation probably plays a stronger role for labile traits that are constantly adjusting to the environment. Additionally, trait plasticity is often related to environmental tolerance, with most studies predicting that higher plasticity leads to broader tolerance — although this depends on how traits map to fitness, given that plasticity can also be maladaptive (Gabriel et al., 2005; Ghalambor et al., 2007; Lande, 2014). In terms of the evolutionary consequences of plasticity, there is general consensus that it either slows or promotes adaptation, yet no clear understanding of the conditions that favour either outcome (Ashander et al., 2016; Chevin & Lande, 2010; Nunney, 2016).

A second theme of the models surveyed concerns population persistence in the face of environmental change, which also involves the evolution of plasticity and tolerance. These models usually incorporate demographic responses to environmental change, given that population size is critical to predictions of persistence *versus* extinction (Gomulkiewicz & Holt, 1995). A good portion of this work sought to identify critical rates of environmental change above which populations can no longer avoid extinction via adaptation (Burger & Lynch, 1995; Chevin et al., 2010; Lynch & Lande, 1993; Osmond & Klausmeier, 2017) or, more generally, the conditions favouring evolutionary rescue, including plasticity and population size (Chevin & Lande, 2010; Gomulkiewicz & Holt, 1995). In this context, genetic variation is also predicted to be important, although it can either favour rescue in highly variable and predictable environments, or have the opposite effect in constant or unpredictable ones (Lande & Shannon, 1996). More recent studies note the importance of frequency-dependent selection, which tends to limit adaptive capacity and persistence in most scenarios of environmental change (Engen et al., 2020; Svensson & Connallon, 2019). Additionally, frequency-dependent selection in temporally fluctuating environments can result in complex and potentially chaotic dynamics, affecting the predictability of evolution and therefore warranting further research (Chevin et al., 2022; Rego-Costa et al., 2018).

Beyond the gaps identified above, we highlight some key areas we believe can move the field forward and improve our ability to predict population responses to climate change. Even though all major components of temporal environmental change (abrupt change in mean, trend, cyclic change, and noise) are relevant in the context of climate change, no study has yet combined them in a unified framework, with or without evolution of plasticity (Figure 3D). Doing so would access key questions about how they contribute to environmental predictability, and their relative impacts on population and evolutionary responses. Further, studies rarely, if ever, attempt to parameterise such models with real or projected time series data to predict such responses into the future. Translating theory from paper to environmental conditions in nature is an important next step in testing and validating predictions (Chevin et al., 2013), with the potential to improve our understanding of biological responses to climate change.

### Predicting the evolution of adaptation and plasticity from components of temporal change

To synthesize theoretical predictions, and assess the relative effects of different components of temporal environmental change on the evolution of adaptation and plasticity, we simulated evolution under different forms of temporal change. First, we developed equations to simulate environmental time series capturing major components of change in Figure 2 (Supporting Information 2A). Next, we applied simulated time series to a quantitative genetic model for the evolution of environmental tolerance, via adaptive changes in the additive genetic breeding value and plasticity of an underlying trait that determines fitness (Supporting Information 2B; Chevin & Lande, 2010; Chevin et al., 2010; Lande, 1976, 2009, 2014). We validated the model by replicating scenarios simulated in earlier studies that retrieved similar predictions (Supporting Information 2C).

We then simulated evolution under different forms of temporal change, using new time series with components of change manipulated relative to a baseline scenario (Supporting Information 2D) and all other parameters held constant. The baseline had an initial mean environment (*ε*_0_) of 5, no trend (*η*_*trend*_ = 0), equal amounts of cyclic change and noise (*η*_*cycle*_ = *η*_*noise*_ = 1), and noise that was uncorrelated or white in colour (*ρ* = 0). To assess the isolated effects of different components on evolution, we simulated a two-fold increase in the initial mean environment (*ε*_0_ = 10), a positive trend (*η*_*trend*_ = 0.001), a four-fold increase in the amplitude of cyclic change (*η*_*cycle*_ = 4), a two-fold increase in noise (*η*_*noise*_ = 2, maintaining a range similar to that of cyclic change), and noise that was positively correlated or reddened in colour (*ρ* = 0.65, approximating *β* = 1). To assess the combined effects of different components on evolution, we simulated pairwise combinations of the changes above. Mean environment was constant unless trend was manipulated.

For different forms of temporal change, we analysed the evolution of tolerance, rates of evolutionary adaptation to track new optima, and the evolution of plasticity relative to environmental predictability (using further simulations detailed below). Data processing and modelling was done in R v4.0.5 (R Core Team, 2021), with code provided on GitHub (https://github.com/CristobalGS/Adaptation-to-TemporalChange).

#### Evolution of environmental tolerance

Predictions exist for evolutionary responses to isolated components of temporal change, but rarely assess responses to them changing in combination, as occurs in nature. Here, we synthesize earlier work on components changing in isolation (diagonals in Figure 4), and further assess their combined effects (off-diagonals in Figure 4). For brevity, we focus on evolved tolerance curves, but present equilibrium breeding values and plasticities used to predict them in Table S3.

**Figure 4.**
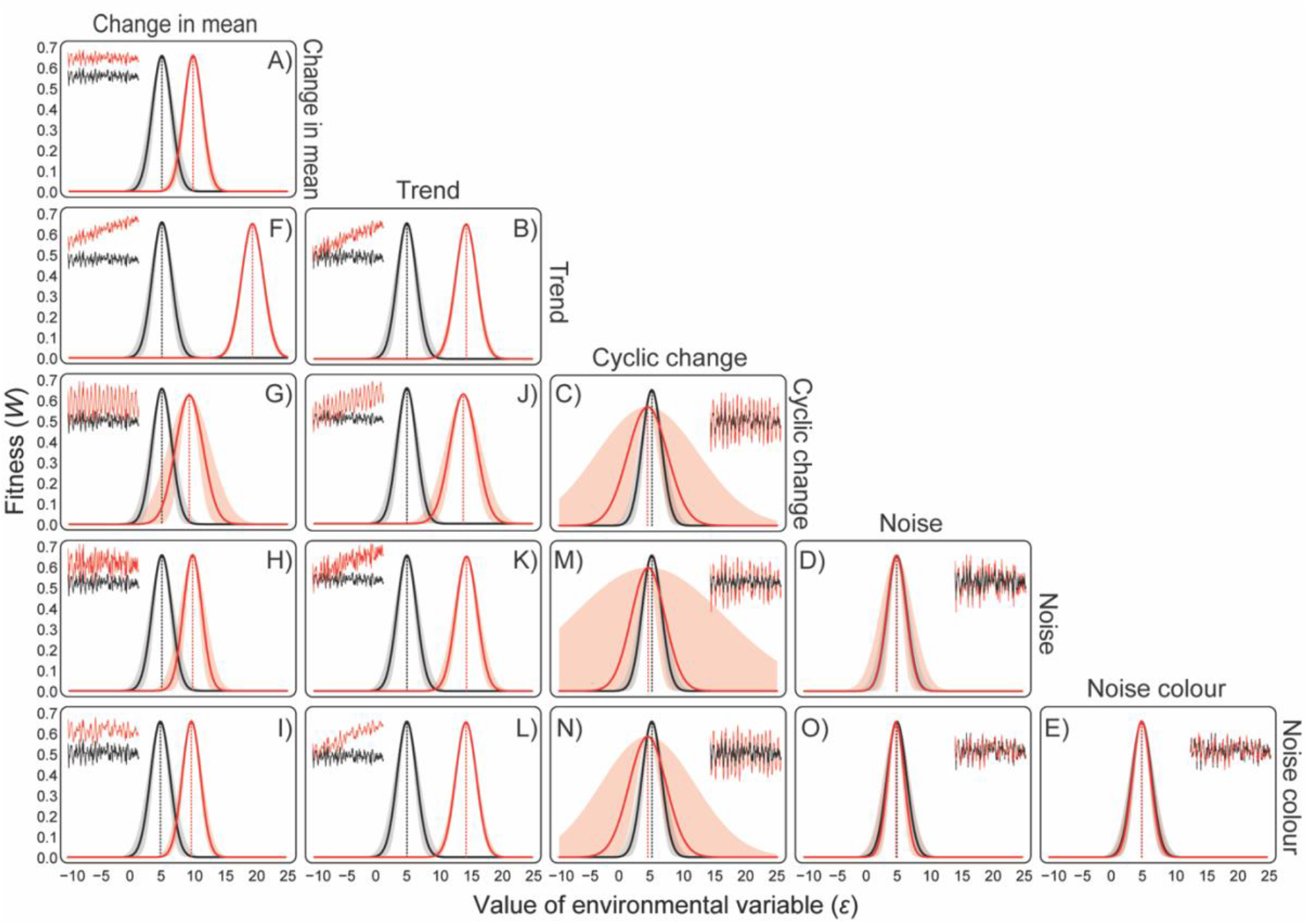
Predicted evolution of environmental tolerance under (A) an abrupt change in the mean environment, (B) a constant rate of change in the mean environment (trend), (C) an increase in the amplitude of cyclic change in the environment, (D) an increase in the amount of stochastic change (noise) in the environment, and (E) an increase in the autocorrelation of stochastic change (noise colour). The evolved tolerance curve for the baseline scenario (Supporting Information 2D) is in black, and curves for alternative scenarios differing in components of environmental change (see main text) are in red. Diagonal plots (A-E) show the effects of components changing in isolation and off-diagonal plots (F-O) show the effects of combined changes. Insets show excerpts of the environmental time series for which evolution was simulated, with the baseline in black and alternative scenarios in red (axes are omitted for clarity but match Figure 2). Curves were predicted using mean breeding values and plasticities at dynamic equilibrium as described in Supporting Information 2B. Shaded areas show variation in tolerance breadths due to variation in plasticities around equilibrium values.

##### Effect of change in mean

The abrupt increase in mean produced higher environmental tolerance relative to the baseline (Figure 4A), with the population shifting the peak of its tolerance curve to match the new environmental optimum. This matched earlier theory (Gomulkiewicz & Holt, 1995) and patterns of directional selection in natural populations (Hoekstra et al., 2001; Kingsolver & Diamond, 2011). Tolerance breadth remained similar to the baseline.

##### Effect of trend

The isolated increase in trend also produced higher tolerance (Figure 4B), but the population now tracked the gradual increase in environmental optimum with a constant lag, as in Lande & Shannon (1996). Tolerance breadth again remained similar to the baseline, though trait plasticity was slightly higher at equilibrium (Table S3).

##### Effect of cyclic change

The isolated increase in cycle amplitude saw plasticity evolve to a higher level relative to the baseline (Figure 4C), increasing both the breadth of environmental tolerance (at the cost of curve height) and the variability in breadth. Similar results were obtained by previous theory (e.g., Ezard et al., 2014) and empirical work (Ketola et al., 2013). Relative to the baseline, moreover, the curve peak matched the mean environment (identical in both scenarios) less accurately.

##### Effect of noise

The isolated increase in noise had little effect beyond increasing the variability in plasticity and hence tolerance breadth (Figure 4D), as expected if temporal change must be predictable for plasticity to evolve (Reed et al., 2010; Tufto, 2000). The isolated increase in noise colour (autocorrelation of noise) also had little effect here (Figure 4E).

##### Combined effects of different components

Most notably, the joint increase in mean and cycle amplitude (Figure 4G) produced higher tolerance, greater tolerance breadth, and greater variability in breadth, but reduced the mean breeding value by ∼64% relative to the isolated increase in mean, and reduced plasticity by ∼62% relative to the isolated increase in cycle amplitude (Table S3). Further, jointly increasing cycle amplitude and noise (Figure 4M) reduced tolerance breadth relative to the isolated increase in amplitude, inferring that stochastic change hinders the effect of cyclic change on plasticity evolution. Other combined changes had similar effects to components changing in isolation.

#### Rates of evolutionary adaptation

To survive environmental change, populations must not only adapt to that change, but adapt fast enough to avoid extinction in the meantime (Bell, 2012; Carlson et al., 2014). Rates of adaptation are therefore key to predicting species’ responses to rapid and complex climate change (Catullo et al., 2019; Garcia et al., 2014; Hoffmann & Sgrò, 2011; Visser, 2008) yet, to our knowledge, no studies have assessed how different components of change can affect them. Here, we analyse the rates at which mean breeding values and plasticities responded to scenarios where abrupt change in mean — alone, and paired with change in other components — imposed directional selection towards new stationary optima. Change in mean and trend (a moving optimum) was excluded given our focus here on adaptation to a new stationary optimum. To do so, we sampled the evolutionary trajectory of each parameter until it approached dynamic equilibrium (∼250-400 generations), smoothed short-term cycles by modelling parameter values as a polynomial function of time, then calculated instantaneous rates of change in values as the tangent slope (or derivative) of the function at each generation.

Adaptation was fastest, and mean breeding value evolved to be highest (matching curve peaks in Figure 4), after an abrupt increase in mean environment alone or with an increase in either component of noise (Figure 5A-B). Jointly increasing mean and cycle amplitude slowed adaptation and produced a lower breeding value (Figure 5A-B), whereas plasticity also evolved slowest, but evolved to be highest, in this scenario compared to all others (Figure 5C-D: note the negative axis in panel D). Since it was the only scenario that slowed adaptation, and the only one in which higher plasticity evolved, the buffering of selection by plasticity could potentially explain these results (Fox et al., 2019; Gibert et al., 2019). Other work has argued, however, that plasticity can aid adaptation to abrupt environmental change — for example, by taking populations closer to their new optima and ‘buying time’ to adapt through genetic assimilation, or revealing cryptic genetic variation for selection to act on (Diamond & Martin, 2021; Fox et al., 2019; Lande, 2009; Schlichting, 2021), although our model does not account for these possibilities. In contrast to cyclic change, changes in noise had little effect on adaptive tracking of new optima here, although this might not hold if noise around new optima was more extreme relative to shifts in optima.

**Figure 5.**
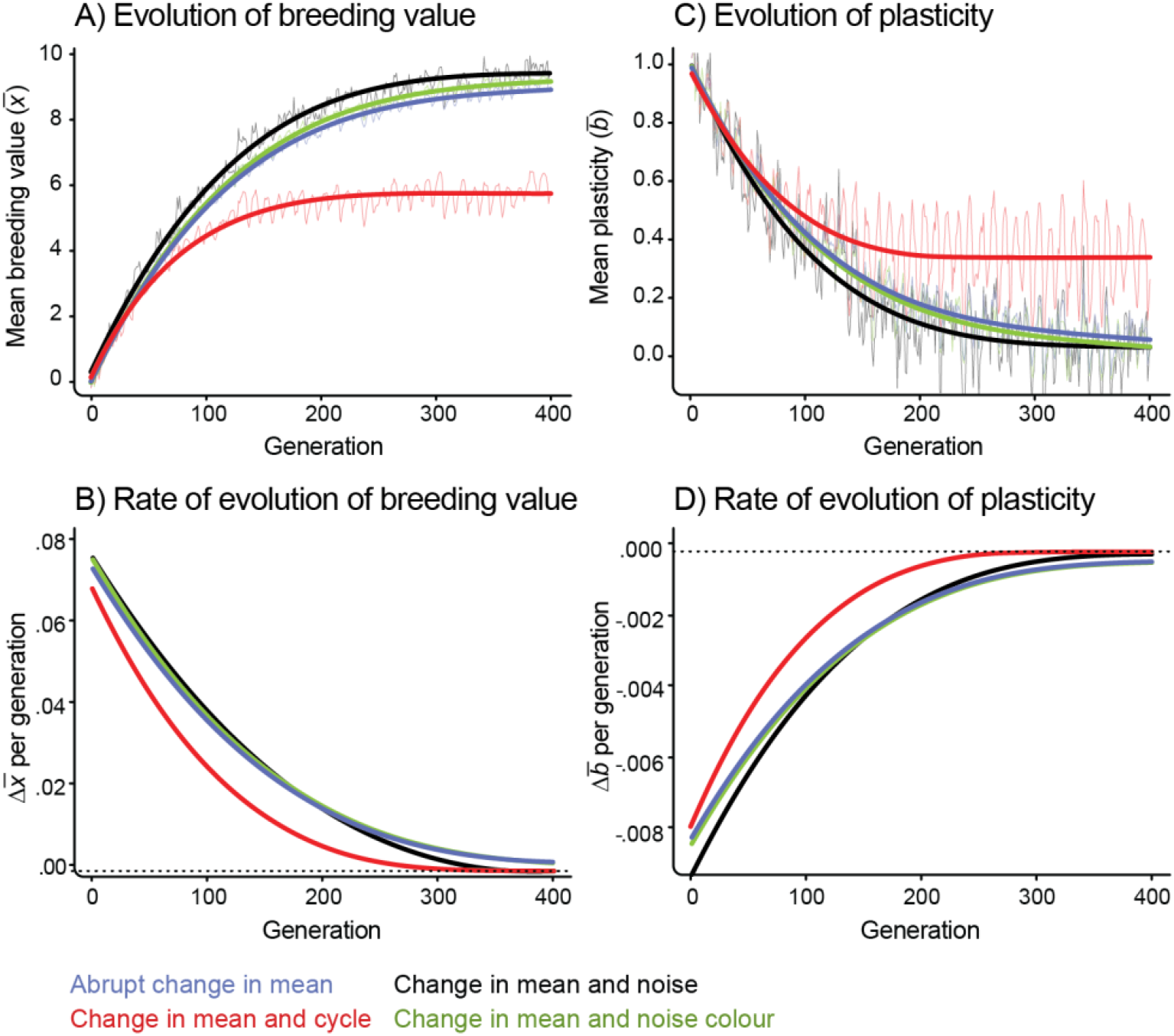
Trajectories and rates of evolution of mean breeding values (A-B) and plasticities (C-D) in environments that impose directional selection towards a new stationary optimum, including an abrupt change in mean (blue), change in mean and cycle amplitude (red), change in mean and noise (black), and change in mean and noise colour (green). Change is relative to the baseline scenario (Supporting Information 2D) and corresponds to the first column of Figure 4, with change in mean and trend (a moving optimum) excluded given our focus here on adaptation to a new stationary optimum. Top panels show raw trajectories (jagged lines) smoothed by regression (smooth lines). Bottom panels show rates calculated from smoothed trajectories. Results are truncated at 400 generations, when trajectories approach equilibrium and rates approach 0 (horizontal dashed lines).

#### Environmental predictability and evolution of plasticity

Since organisms need reliable cues to respond to, environmental predictability is considered necessary for adaptive plasticity to evolve (Bitter et al., 2021; Lande, 2014; Reed et al., 2010). Less recognised, however, is that different components of temporal change can shape predictability of different forms, with potentially different outcomes (Botero et al., 2015; Ezard et al., 2014; Marshall & Burgess, 2015). Here, we analyse the evolution of plasticity in environments with the same mean and range, but dominated by either cyclic change (*η*_*cycle*_ = 4, *η*_*noise*_ = 1) or noise (*η*_*cycle*_ = 1, *η*_*noise*_ = 2), and differing in the autocorrelation or colour of noise (0 ≤ *ρ* ≤ 0.9). We manipulated noise colour because it is often used as a proxy for predictability (Ruokolainen et al., 2009; Vasseur & Yodzis, 2004), but also explored predictability in terms of the time lag between development and selection, manipulating generation time (*T*). To simplify, we here focused on three cases that cover most dynamics and emerging patterns based on preliminary tests: (i) adults experienced a complete environmental cycle before selection (*T* = 11, as in simulations above), (ii) only a partial cycle before selection (*T* = 2), or (iii) no environmental change before selection (*T* = 1). In an attempt to unify different forms of predictability, we calculated the autocorrelations of time series with all components combined (using R’s *pacf* function) and the resulting correlations between environments of development and selection. Our analysis provides new insights into the complex effects of predictability on the evolution of plasticity.

When cyclic change dominated (Figure 6A), the autocorrelation of the time series ranged from 0.7 to 0.8 and was perfectly correlated with the autocorrelation of noise alone (with slope relating the first in response to the second ∼0.1). In this case, plasticity at evolutionary equilibrium depended little on noise colour, changing mainly as environmental change within generations altered the correlation between the environment that induced plasticity in the trait and the environment of selection on the trait. Plasticity evolved to its highest levels in the absence of within-generation change (*T =* 1), when those environments were the same and hence perfectly correlated. We modelled this scenario by eliminating the lag between development and selection, which could also approximate the case of a labile trait adjusting so rapidly to environmental cues that development and selection occur simultaneously and continuously in time (e.g., Lande, 2014). Evolved plasticity then declined as within-generation change (*T* > 1) decoupled the environments of development and selection, reaching its lowest levels when adults encountered only a partial cycle of change before selection (*T =* 2) and those environments were most different. Note that although not shown here, the correlation between the environments of development and selection continued to decrease down to negative values between 2 < *T* < 11, as this further increased the mismatch between those environments, thus leading to a reversal in the slope of the evolved reaction norm.

**Figure 6.**
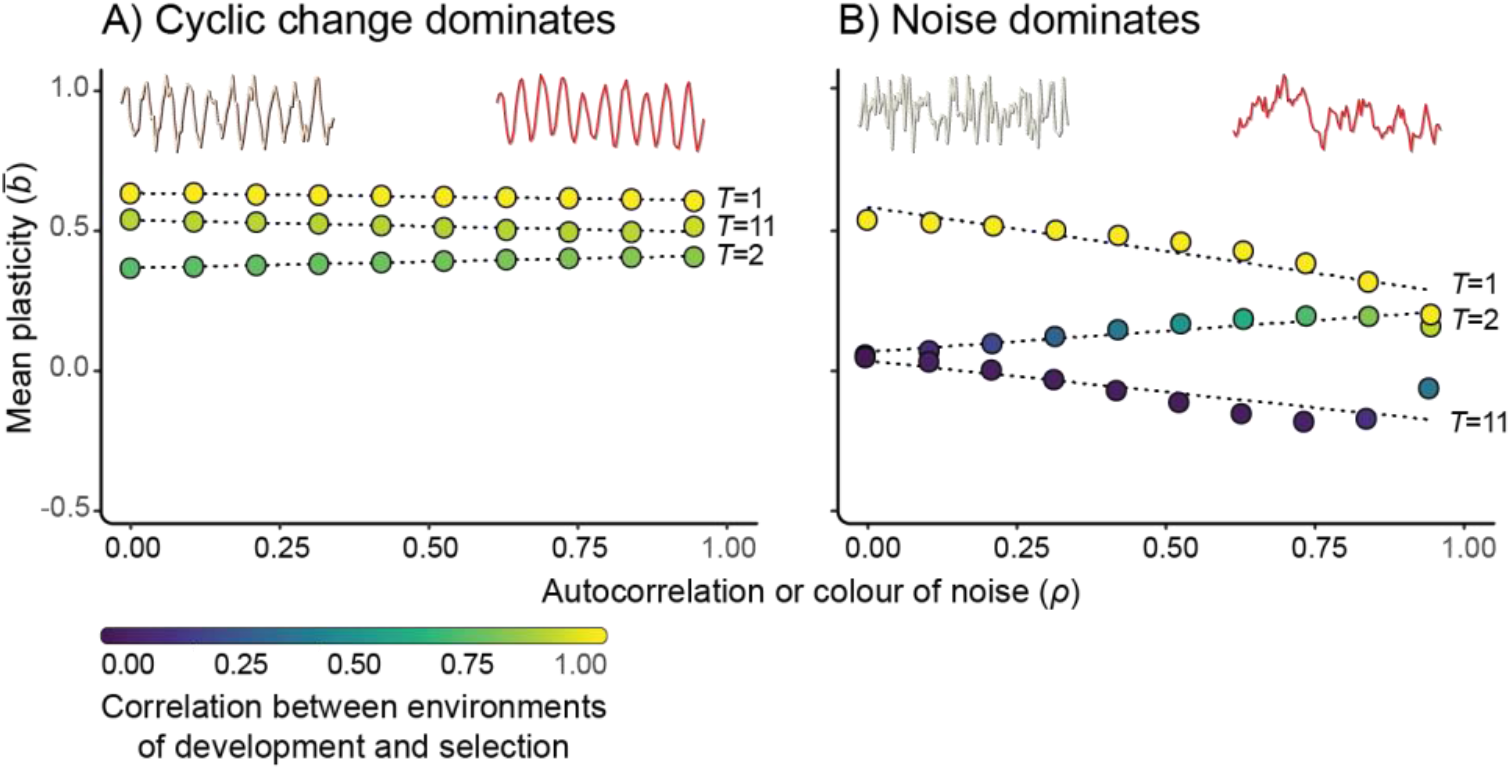
Evolution of plasticity under different forms of predictability in environments. A) dominated by cyclic change or B) dominated by noise. Noise ranges from uncorrelated or white in colour (*ρ* = 0) to positively correlated or reddened in colour (*ρ* = 0.9). Each point is an individual simulation coloured by the correlation between environments of development and selection. Sets of simulations differ in generation time, exposing adults to either a complete cycle of environmental change before selection (*T* = 11), only part of a cycle before selection (*T* = 2), or no change before selection (*T* = 1). Dashed lines show trends for different generation times. Insets show excerpts of the environmental time series for which evolution was simulated, with whiter noise on the left and reddened noise on the right (axes are omitted for clarity but match those for Figure 2).

When noise dominated (Figure 6B), the autocorrelation of the time series ranged from 0.1 to 0.9 and was perfectly correlated with the autocorrelation of noise alone (the slope relating them was ∼0.9). In this case, equilibrium plasticity was still highest in the absence of environmental change within generations, but depended jointly on generation time and noise colour. In the absence of within-generation change (*T =* 1, when environments of development and selection were the same), more autocorrelated, reddened noise led to declines in plasticity. Although this seems to contrast with the common theoretical prediction (e.g., Gavrilets & Scheiner, 1993; Lande, 2009) it is actually a scenario that is not usually modelled. In this particular case, a higher environmental autocorrelation combined with simultaneous development and selection may allow the mean phenotype to track the optimum phenotype more closely by evolution of the mean breeding value (i.e., reduced stochastic load that increases adaptive tracking; Ashander et al., 2016; Lande & Shannon, 1996), thus favouring less plasticity (given a cost to it) via genetic assimilation (Lande, 2009). In other scenarios, however, higher autocorrelation increased plasticity (as expected) by offsetting the degree to which within-generation change decoupled the environments of development and selection. Those environments differed most when adults encountered a longer time lag before selection (*T =* 11), but plasticity evolved to similar levels in this scenario — albeit with negative values indicating a reversal in reaction norm slope — as it did when adults encountered a shorter time lag before selection (*T =* 2). Although we did not model it, negative environmental autocorrelation over a developmental time lag can select a reaction norm with slope of opposite sign to that of the optimum phenotype as a function of the environment (Lande, 2009), and it is possible that the interplay of noise colour and lag produced a similar dynamic here.

Unifying different forms of environmental predictability highlights the correlation between environments of development and selection as an integrated measure that best explains the evolution of plasticity (Bitter et al., 2021; Gavrilets & Scheiner, 1993; Lande, 2009). Nevertheless, it did not always map to autocorrelation in the time series or the lag between development and selection, nor did it always explain plasticity. The effect of lag depended on cycle period relative to lag length in series dominated by cyclic change, where autocorrelation was always high regardless of noise colour, but lag length alone in series dominated by noise, where autocorrelation equated to noise colour. This emphasises that life history may critically shape how plasticity evolves in response to different components of temporal change — for example, we might expect it to respond more to noise colour in short-lived microbes that do not experience environmental cycles (e.g., Leung et al., 2020), but to? seasonality relative to generation time in longer-lived species (reviewed by Williams et al., 2017). However, plasticity did not always respond to predictability as expected when it was induced and selected simultaneously, arguing that theory tailored for developmentally plastic traits may not hold for those adjusted continually through life (see also Rescan et al., 2022). Since fewer models have explored the dynamics of labile traits (Beaman et al., 2016; Lande, 2014; see also review section above), this points to a need to better understand the role of predictability in their evolution (Rescan et al., 2022).

#### Case Study: Predicting thermal adaptation and plasticity in a global marine hotspot

Temperature will almost certainly drive adaptation, given its pervasive biological effects (Clarke, 2017; Kingsolver, 2009). Predicting thermal adaptation in nature may therefore be crucial in predicting species’ responses to ongoing climate change (Hoffmann & Sgrò, 2011). We illustrate the utility of our model in this regard by applying it to a case study of a global marine hotspot in southern Australia (Costello et al., 2022; Hobday & Pecl, 2014). Marine hotspots such as this not only harbour exceptional biodiversity, but have warmed more and faster than terrestrial hotspots (Costello et al., 2022), and biodiversity in them is especially prone to redistribution, given that marine species track climate velocities more closely than terrestrial species (Lenoir et al., 2020; Sunday et al., 2015). To predict thermal adaptation and plasticity across the hotspot, we parameterised our model with daily time series of sea surface temperature for several locations across the hotspot that differed mostly in mean, cyclic change (measured by annual range), and noise colour (see details in Box 1 and Box 2).

Our simulations predict that thermal environments across this region favour local adaptation, with populations differing in both the positions and (to less extent) breadths of their thermal tolerance curves. These predictions — and the parameter values they rely on — could be validated empirically, for example, by exploring patterns of adaptation and plasticity using common gardens or reciprocal transplants (Blanquart et al., 2013), estimates of selection and heritability (Shaw & Etterson, 2012), and genomic tools such as allele-environment associations and expression analyses (Mallard et al., 2020; Rellstab et al., 2015). Moreover, that locations differed less in plasticity (and hence tolerance breadth) than expected from differences in thermal range highlights that reconciling model predictions with expected outcomes is not straightforward, and warrants careful consideration of whether extrinsic cues for plasticity, as well as intrinsic costs and limits to its evolution (DeWitt et al., 1998) are adequately captured, Nevertheless, our model generates testable null expectations, based on real-world environments, that can guide future research and paths for validation.

#### Concluding remarks

Environmental change is complex, and our ability to predict evolutionary responses to it is still poor (Rescan et al., 2022). In particular, there is growing recognition of the need to better account for the complexity of temporal environmental change in biological predictions (Bates et al., 2018; Dillon et al., 2016; Helmuth et al., 2014; Vinton et al., 2022). Our brief review of theory on adaptation to temporal environmental change identified two key gaps. First, models rarely integrate multiple components of temporal change in a unified framework to study their relative and combined effects on adaptation, or the plasticity underpinning it. Second, only in rare occasions have theoretical studies attempted to directly translate their predictions to patterns of temporal change in natural environments. Biologists and environmental managers have much to gain from progress in both areas. To address those gaps, we used existing theory to develop a quantitative genetic model for the evolution of adaptation and plasticity in response to multiple, major components of temporal change. First, we retrieved and synthesised previous predictions, then generated new insights into the effects of different types of change on evolutionary outcomes, including plasticity in relation to environmental predictability. Second, we applied our model to a case study of temporal change in temperature across the southern Australian hotspot, to illustrate how it may be used to predict adaptation and plasticity, and guide efforts to better understand them, in priority regions for conservation where biodiversity may be particularly sensitive to climate change.

We aimed to make our model general and able to be tailored to any system with environmental time series available. For example, it could be used with climatic variables scaled to different temporal resolutions to match ecologically- and evolutionarily-relevant scales of climate change, or with climate projections to predict adaptation in future climates. Nevertheless, we acknowledge that all models are compromises of generality, precision, and realism (Levins, 1966), and ours is no exception. Evolution may of course depend on other factors, including life-history, population structure and gene flow, and genetic architecture, whose effects were not explored here (but see, e.g., Agrawal & Stinchcombe, 2009; Hadfield, 2016; Pease et al., 1989). For example, local adaptation might not evolve if gene flow among populations is high (Hadfield, 2016; Scheiner, 1998; Tigano & Friesen, 2016), if populations lack sufficient genetic variation for adaptation (Hoffmann & Merilä, 1999; Lande & Shannon, 1996), or if genetic correlations inhibit adaptation (Chevin, 2013; Lande, 1979). Information on populations’ physiological limits (Catullo et al., 2015), adaptive capacities (Bush et al., 2016), and genetic structures (Macdonald et al., 2018; Prunier & Blanchet, 2018) would all be useful to improve model predictions and their geographic context. Much of this information, if available, could already be imputed in our model, or incorporated by slightly modifying it. Future research could also relax the shapes of tolerance curves and/or reaction norms, and consider labile traits (Lande, 2014) or multiple correlated traits (Lande & Arnold, 1983) to improve predictions. Our approach is well-established, broadly applicable, and offers an accessible framework for expanding understanding of evolutionary adaptation in the field, with scope to enhance the management and conservation of biological systems.

## Acknowledgements

We specially thank Tim Connallon for his valuable help developing the mathematical model, the literature review, and general advice on earlier versions of the manuscript. This research was supported by The Holsworth Wildlife Research Endowment & The Ecological Society of Australia, awarded to CG, and by grants awarded under the Australian Research Council’s Discovery Scheme to KM and KH.

## Conflict of interest

The authors have no conflict of interest to declare.

## Data availability statement

The reproducible R scripts with simulations, analyses, and visualisations that supports the findings of this study will be available on GitHub (https://github.com/CristobalGS/Adaptation-to-TemporalChange) upon publication.

## Box 1. Temporal change in temperature across the southern Australian marine hotspot

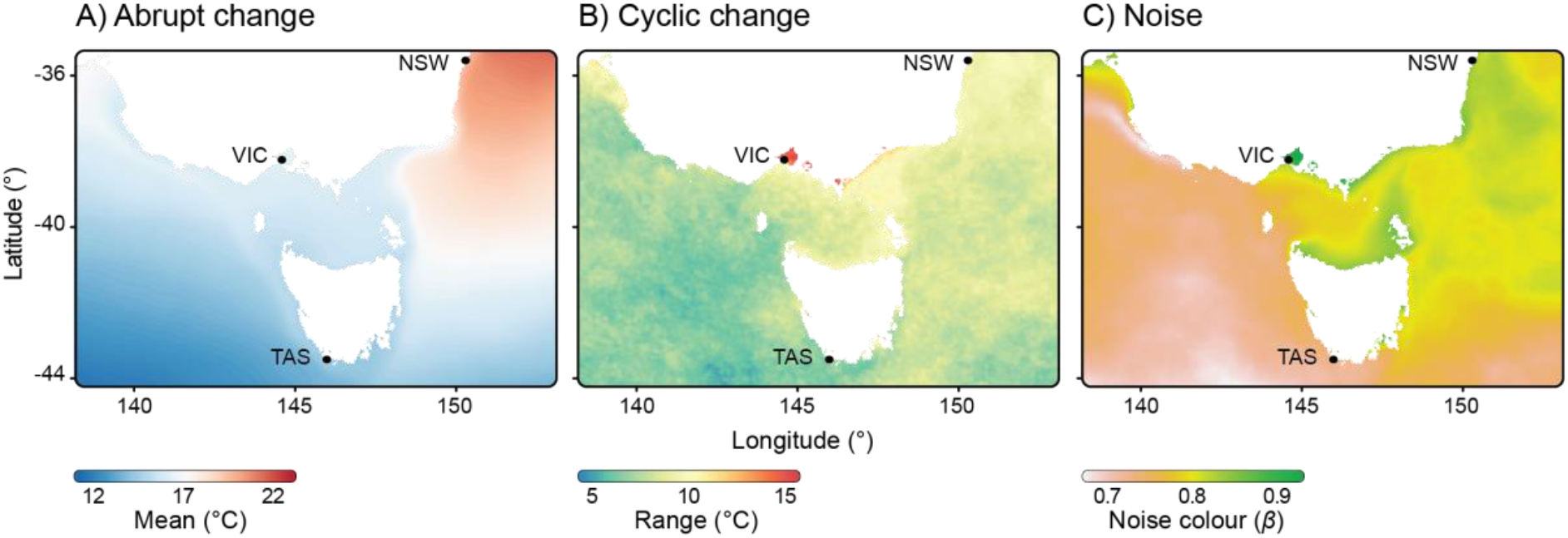

Southern Australia is a global hotspot of marine biodiversity (Costello et al., 2022) and warming much faster than the global average rate (Hobday & Pecl, 2014). In the hotspot, strongly seasonal currents flowing south from the tropics and east from the Great Australian Bight meet subantarctic water in Bass Strait, causing temporal change in temperature to vary regionally and with localised features like bays and upwellings (Wijffels et al., 2018). To apply our model, we obtained high-resolution (1 km x 1 km grid) daily observations of sea surface temperature from 2011 to 2018 (www.ghrsst.org). First, to identify geographic differences in temperature change, we estimated the mean, range (a measure of cyclic change), and autocorrelation of stochastic change (noise colour) for each grid cell of interest, and mapped components (panels A–C). Other components showed little change across the hotspot and were therefore omitted here for brevity. Mean and cyclic change were estimated as averages of the bioclimatic variables BIO1 (annual mean temperature) and BIO7 (temperature annual range, which approximates seasonal differences in temperature) in the WorldClim scheme (see Fick & Hijmans, 2017). Noise colour was estimated as *β* (see Figure 2 and Vasseur & Yodzis, 2004). Second, to parameterise the model with real temperature data and based on geographic variation visible in maps, raw time series were extracted for three locations (dots in maps, labelled by the state they are in) – southwestern Tasmania (TAS), Port Phillip Bay in Victoria (VIC), and southern New South Wales (NSW) – which were then fed to model simulations.

## Box 2. Predicted thermal adaptation and plasticity in the southern Australian marine hotspot

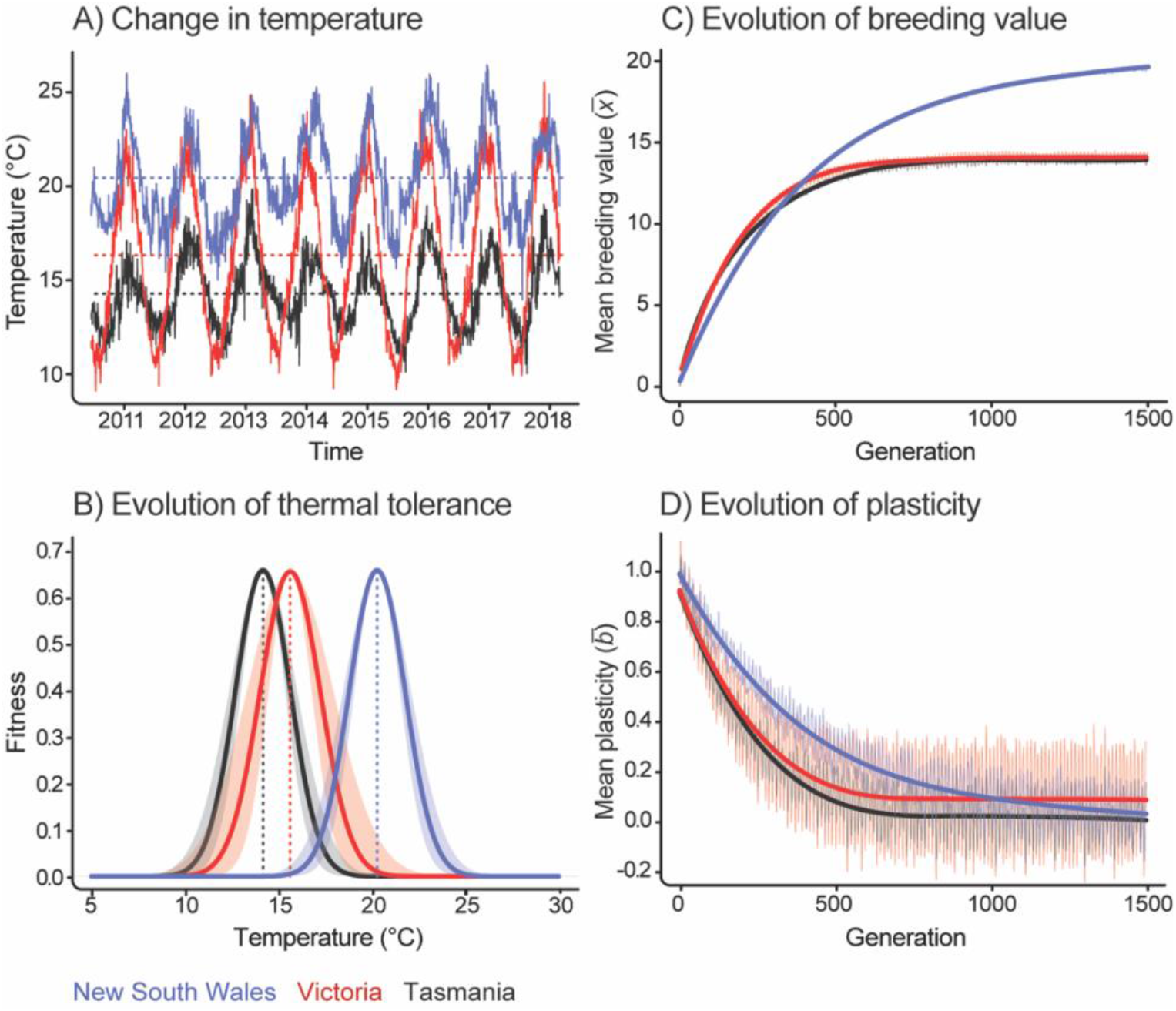

To predict thermal adaptation and plasticity across the southern Australian hotspot, we parameterised our model with daily time series of sea surface temperature for several locations that differed mostly in mean, cyclic change (measured by annual range) and noise colour (Box 1). We set generation time (*T*) to 30 days to simulate a common life cycle duration in marine invertebrates (Levin & Bridges, 1995). Other parameters (including trait variance and heritability, strength of stabilizing selection, and environmental sensitivity of selection) matched the baseline scenario (Supporting Information 2D). In the absence of empirical estimates, we assumed that they were uniform across locations, and confirmed that altering them uniformly did not qualitatively change our results (e.g., stronger stabilizing selection reduced tolerance breadth overall, but did not change how breadths compared across locations). Since outcomes did not reach dynamic equilibrium during the time series, and trend for the period (2011–2018) was near-zero at each location, we repeated series 20 times to simulate ∼1900 generations in total.

Time series for selected locations (panel A, with horizontal lines showing trends in means) translated to predictions of geographic variation in thermal adaptation at evolutionary equilibrium (panel B). Tolerance curves differed predominantly in the positions of optima (vertical lines), predicting higher thermal tolerance through evolution of higher breeding values in locations (such as New South Wales) with higher mean temperatures (panel C). Evolved plasticities also varied geographically, but in more complex ways (panel D). Plasticity at equilibrium was highest in Victoria and lowest in Tasmania, but the difference was minor and translated to little difference in tolerance breadth among locations (panel B). Rather, Victoria’s greater thermal range translated to greater variability in tolerance breadth (shaded area in panel B) as plasticity fluctuated more around its equilibrium value (panel D). For much of the non-equilibrium evolution, however, conditions favoured considerably higher plasticity (and hence broader tolerance) in New South Wales than in southern locations (panel D). Raw trajectories (jagged lines) of breeding values and plasticities are smoothed by regression (smooth lines) and truncated at equilibrium (see Table S4 for equilibrium values underpinning curves in panel B).

## Notes

https://github.com/CristobalGS/Adaptation-to-TemporalChange

### Competing Interest Statement

The authors have declared no competing interest.

