## Supporting information for "Predicting the evolution of adaptation and plasticity from temporal environmental change"

**1. Theoretical studies of adaptation to temporal environmental change.**

**Table S1.** Targeted searches for relevant theoretical studies. Studies were initially scanned based on title and/or abstract and then scanned for content if identified as relevant. Studies were included if they explicitly modelled evolution of adaptation (focussing on trait(s) or fitness directly) in response to temporal environmental change. Therefore, experimental studies, studies that modelled demographic but not evolutionary responses, and studies focusing exclusively on spatial environmental change, were excluded. Searches were conducted in April 2022.

**Steps in searches**

Identified studies based on previous knowledge

Checked all studies cited in Chevin et al. 2010

Checked all studies by R. Lande (Google Scholar profile)

Checked all studies by L-M Chevin (Google Scholar profile)

Checked all studies cited in Lande 2014

Checked all studies cited in Lande 2019

Checked all studies cited in Chevin et al. 2022

Checked all studies cited in Kopp and Matuszewski 2014

Checked all studies cited in Coulson et al. 2017

Searched Google Scholar for term ‘phenotypic evolution * environmental change’ and checked first 100 articles (sorted by descending relevance)

Searched Google Scholar for term ‘tolerance * evolution * environmental change’ and checked first 100 articles (sorted by descending relevance)

**Table S2.** Theoretical studies included in targeted literature review on adaptation to temporal environmental change. For inclusion criteria see Table S1. Raw data extracted from them (summarised in Figure 3) can be provided upon request.

**2. Evolution of adaptation and plasticity under different forms of temporal environmental change**

**A) Modelling environmental change.** We modelled components of temporal change in Figure 2 as a sum of independent terms. The value of the environmental variable ($\varepsilon$) at any given time (*t*) was given by

$\varepsilon_{t}=\varepsilon_{0}+\eta_{trend}t+\eta_{cycle}\sin\left( 2\pi t/P \right)+\eta_{noise}\zeta$	(eq. 1),

where $\varepsilon_{0}$ is the intercept or initial mean value, $\eta_{trend}, \eta_{cycle}, and \eta_{noise}$are scaling constants reflecting the relative importance of trend $(\eta_{trend}t$), cyclic change ($\eta_{cycle}\sin\left( 2\pi t/P \right),$ defining a sine wave with amplitude $\eta_{cycle}$ and period *P*), and noise ($\eta_{noise}\zeta$; see below) in the variable. Noise colour (the autocorrelation of noise) was modelled as

$\zeta_{t+1}= {\rho\zeta}_{t}+ \gamma_{t}\sqrt{1-\rho^{2}}$	(eq. 2),

where $\rho$ is the autocorrelation coefficient, and $\gamma_{t}$ is a standard normal random variable (Ruokolainen et al., 2009). Hence, $\zeta$ can range from uncorrelated white noise when $\rho=0$ to positively correlated reddened noise when $\rho>0$, without changing its variance (see Figure 2).

**B) Modelling evolution of adaptation and plasticity.** We modelled an individual’s expression of a quantitative trait ($z$) as the sum of genetics, phenotypic plasticity, and random residual variation as follows

$z=x+b\varepsilon_{t}+y$	(eq. 3),

where $x$ is the additive genetic breeding value for the trait (the elevation of the linear reaction norm relating it to $\varepsilon_{t}$, the environmental variable in eq. 1), $b$ is plasticity in the trait (the slope of the reaction norm), and $y$ is residual variation. Terms $x$, $b$, and $y$ are independent and normally distributed random variables, $x\sim N(\bar{x}, \sigma_{x}^{2})$, $b\sim N(\bar{b}, \sigma_{b}^{2})$, $y\sim N(\bar{y}=0, \sigma_{y}^{2}=0.1)$, where overbars denote means and $\sigma^{2}$ denote variances.

The trait shows developmental plasticity, so that its expression is determined by environmental cues at the start of the life cycle and remains fixed thereafter (Nettle & Bateson, 2015). For simplicity, and to explore the effect of a lag between the environments of development and selection on the trait, we modelled selection operating at the end of life via fertility, as in semelparous organisms (e.g., Young & Augspurger, 1991). Selection could alternatively be modelled operating throughout the lifecycle (as in Cotto & Chevin, 2020), which could be explored in future work. We assumed that generations are discrete and non-overlapping, and expressed generation time ($T$) in arbitrary time-steps so that populations experience multiple environmental states (modelled by eq. 1) if *T* > 1. Therefore, in each generation *G*, trait expression is determined by the environment at the start of life ($\varepsilon_{\left( G-1 \right)T+1}$), selection occurs in the environment at the end of life ${(\varepsilon}_{GT})$, and if $T=1 ($no environmental change within generations), the environments of development and selection are the same ($\varepsilon_{\left( G-1 \right)T+1}= \varepsilon_{GT}$). We further assumed that all individuals in the population experience the same environment, that population size is infinite and constant (no drift), and that selection is both frequency and density independent.

We allowed evolution by natural selection to change the trait’s mean breeding value ($\bar{x}$) and mean plasticity ($\bar{b}$) in the population. The variances for *x* and *b* ($\sigma_{x}^{2}$ and $\sigma_{b}^{2}$), as well as for the residual effects ($y$), were assumed to be constant. Given the states of $\bar{x}$, $\bar{b}$, and $\varepsilon_{\left( G-1 \right)T+1}$ in each generation, the trait is normally distributed with mean and variance of

$\bar{z}=\bar{x}+\bar{b}\varepsilon_{\left( G-1 \right)T+1}$	(eq. 4a)

$\sigma_{z}^{2}=\sigma_{x}^{2}+\varepsilon_{\left( G-1 \right)T+1}^{2}\sigma_{b}^{2}+\sigma_{y}^{2}$	(eq. 4b).

Fitness in each generation is then a Gaussian function of trait expression as follows

$W\left( z \right)=W_{\max}\exp\left( -\frac{\left( \theta-z \right)^{2}}{2\omega_{z}^{2}}-\frac{b^{2}}{2\omega_{b}^{2}} \right)$	(eq. 5),

where $\theta$ is the trait optimum, $W_{\max}$ is the maximum individual fitness, ${1/\omega}_{z}^{2}$ is the strength of stabilizing selection on the trait, and ${1/\omega}_{b}^{2}$ is the cost of plasticity in the trait (modelled as the strength of stabilizing selection on the reaction norm slope; Chevin et al., 2010; Lande, 2014). The trait optimum is a linear function of the environment at the time of selection, $\theta=A+B\varepsilon_{GT}$, where $A$ is the optimum at $\varepsilon_{0}=0$ (the initial or reference environment) and $B$ is linear change in the optimum with environmental change (known as the environmental sensitivity of selection; Chevin et al., 2010).

Following Chevin & Lande (2010), we assumed that selection in each generation acts first on the trait’s plasticity and then on its mean. After selection on plasticity, the distribution of $b$ is normal with mean and variance of

$\bar{b}^{*}=\frac{\bar{b}\omega_{b}^{2}}{\omega_{b}^{2}+\sigma_{b}^{2}}$	(eq. 6a)

$\sigma_{b}^{2*}=\frac{\sigma_{b}^{2}\omega_{b}^{2}}{\omega_{b}^{2}+\sigma_{b}^{2}}$	(eq. 6b)

(e.g., Lande, 1976; Svensson & Connallon, 2019), so that the trait distribution is normal with mean and variance of

$\bar{z}^{*}=\bar{x}+\bar{b}^{*}\varepsilon_{\left( G-1 \right)T+1}$	(eq. 7a)

$\sigma_{z}^{2*}=\sigma_{x}^{2}+\varepsilon_{\left( G-1 \right)T+1}^{2}\sigma_{b}^{2*}+\sigma_{y}^{2}$	(eq. 7b).

Mean lifetime fitness of the population then becomes

$\bar{W}=W_{\max}\sqrt{\frac{\omega_{b}^{2}\omega_{z}^{2}}{\left( \omega_{b}^{2}+\sigma_{b}^{2} \right)\left( \omega_{z}^{2}+\sigma_{z}^{2*} \right)}}\exp\left( -\frac{\bar{b}^{2}}{2\omega_{b}^{2}+2\sigma_{b}^{2}}-\frac{\left( \theta-\bar{z}^{*} \right)^{2}}{2\omega_{z}^{2}+2\sigma_{z}^{2*}} \right)$	(eq. 8),

and the evolution of the mean breeding value and mean plasticity per generation is given by

$\Delta\bar{x}=\sigma_{x}^{2}\frac{\partial\ln\left( \bar{W} \right)}{\partial\bar{x}}$	(eq. 9a)

and

$\Delta\bar{b}=\sigma_{b}^{2}h_{b}^{2}\frac{\partial\ln\left( \bar{W} \right)}{\partial\bar{b}}$	(eq. 9b),

where $h_{b}^{2}$ is the heritability of plasticity (Lande, 1976).

Finally, substituting $\theta=A+B\varepsilon_{GT}$ and eq. 4a into eq. 8 gives fitness as a function of $\bar{x}$, $\bar{b}$, and $\varepsilon$ as follows

$\bar{W} \left( \bar{x}, \bar{b}, \varepsilon\right)=W_{\max}\sqrt{\frac{\omega_{b}^{2}\omega_{z}^{2}}{\left( \omega_{b}^{2}+\sigma_{b}^{2} \right)\left( \omega_{z}^{2}+\sigma_{z}^{2} \right)}}\exp\left( -\frac{\bar{b}^{2}}{2\omega_{b}^{2}+2\sigma_{b}^{2}}-\frac{\left( A+B\varepsilon_{GT}-\bar{x}-\bar{b}\varepsilon_{1} \right)^{2}}{2\omega_{z}^{2}+2\sigma_{z}^{2}} \right)$	(eq. 10).

We then predicted the evolved environmental tolerance curve (per Figure 1) by using the evolved breeding value ($\bar{x})$, and plasticity ($\bar{b}$) to estimate fitness across an arbitrary environmental range ($\varepsilon$). Breeding value determined curve position along the range (equivalent to $\varepsilon_{opt}$ in Figure 1), plasticity determined curve breadth (with higher plasticity leading to higher breadth, per eq. 10 and Figure 1), and a broader curve entailed a lower peak (maximum fitness) due to the cost of plasticity (Figure 1).

**C) Model validation.** To validate our model, we simulated scenarios from earlier studies and compared our outcomes against those previous predictions. As per Lande & Shannon (1996), when setting plasticity to zero and not allowing it to evolve ($\bar{b}=0, \sigma_{b}^{2}=0$), the mean phenotype evolved rapidly towards a new fixed optimum, or tracked an optimum moving at a constant rate with a constant lag at equilibrium. In the latter case, the observed lag (difference between optimum and trait value) closely matched the expected lag ($\eta_{t}\omega_{z}^{2}/\sigma_{x}^{2}$). When setting initial plasticity to zero but allowing it to evolve ($\bar{b}=0, \sigma_{b}^{2}=0.045$), our model retrieved a similar evolutionary dynamic to Lande (2009), in which adaptation to an abrupt, extreme change in environment first occurs by a rapid transient increase in plasticity, which slowly declines to a final equilibrium value (> 0) as the breeding value slowly approaches the new optimum by genetic assimilation.

**D) Baseline scenario.** As a baseline for comparison, we simulated 10,000 time-steps (e.g., days, months, or years) of evolution for a population that was optimally adapted to its environment initially. Initial mean breeding value was $\bar{x}_{0}=0$, and initial mean plasticity was $\bar{b}_{0}=1$. To simplify the evolutionary analysis, and following Lande (2014), we assumed that the breeding value of the optimum phenotype in the reference environment was $A=0$ and its optimum plasticity was $B=1$, so that mean plasticity relative to its optimum equals mean plasticity ($\bar{b}/B=\bar{b}$; as in Ashander et al., 2016). The initial mean environment was five ($\varepsilon_{0}=0)5$), with no trend ($\eta_{trend}=0$) and equal amounts of cyclic change and noise ($\eta_{cycle}=\eta_{noise}=1$). Cycle period was $P=10$ (10 time-steps per cycle), and noise colour was white (no autocorrelation, $\rho=0$) (Figure S1A). Generation time was $T=11,$ so that each generation experienced a complete cycle of environmental change before selection (as in Lande, 2019). Other parameters had values of $\sigma_{x}^{2}=1$, $\sigma_{b}^{2}=0.1$, $\omega_{b}^{2}=1$, $\omega_{z}^{2}=1$, $y=0$, $\sigma_{y}^{2}=0.1$, $W_{\max}=1$, $h_{b}^{2}=0.5$ (higher and lower costs of plasticity, $\omega_{b}^{2}$, were explored, but led to lower or higher evolved plasticity without qualitatively changing results).$\varepsilon_{0}$

We then tracked the evolution of mean breeding value ($\bar{x}$) and mean plasticity ($\bar{b}$) across generations under this baseline environmental scenario, and predicted the mean tolerance curve (along with its range as a measure of variability) using equilibrium values from generation 400 onwards. At this point, the mean breeding value (Figure S1B) and mean plasticity (Figure S1C) reached dynamic equilibrium (as in Chevin & Lande, 2015), fluctuating stably in ways that made the evolved reaction norm vary in slope (Figure S1D, shaded area), and the evolved tolerance curve vary in both breadth (Figure E, shaded area) and position (shown together in Figure S2). For simplicity, and given variation in curve position did not qualitatively change our results, we here focus on variation in breadth only.

**E) References**. **Ashander et al.** (2016) *Proc R Soc B Biol Sci* 283:20161690. **Chevin & Lande** (2010) *Evolution* 64:1143. **Chevin & Lande** (2015) *Evolution* 69:2767. **Chevin et al.** (2010) *PLoS Biol* 8:e1000357. **Cotto & Chevin** (2020) *Theor Popul Biol* 134:119. **Lande** (1976) *Evolution* 30:314. **Lande** (2009) *J Evol Biol* 22:1435. **Lande** (2014) *J Evol Biol* 27:866. **Lande** (2019) *Proc Natl Acad* Sci 116:11361. **Lande & Shannon** (1996) *Evolution* 50:434. **Nettle & Bateson** (2015) *Proc R Soc B Biol Sci* 282:20151005. **Ruokolainen et al.** (2009) *Trends Ecol Evol* 24:555. **Svensson & Connallon** (2019) *Evol Appl* 12:1243. **Young & Augspurger** (1991) *Trends Ecol Evol* 6:285.

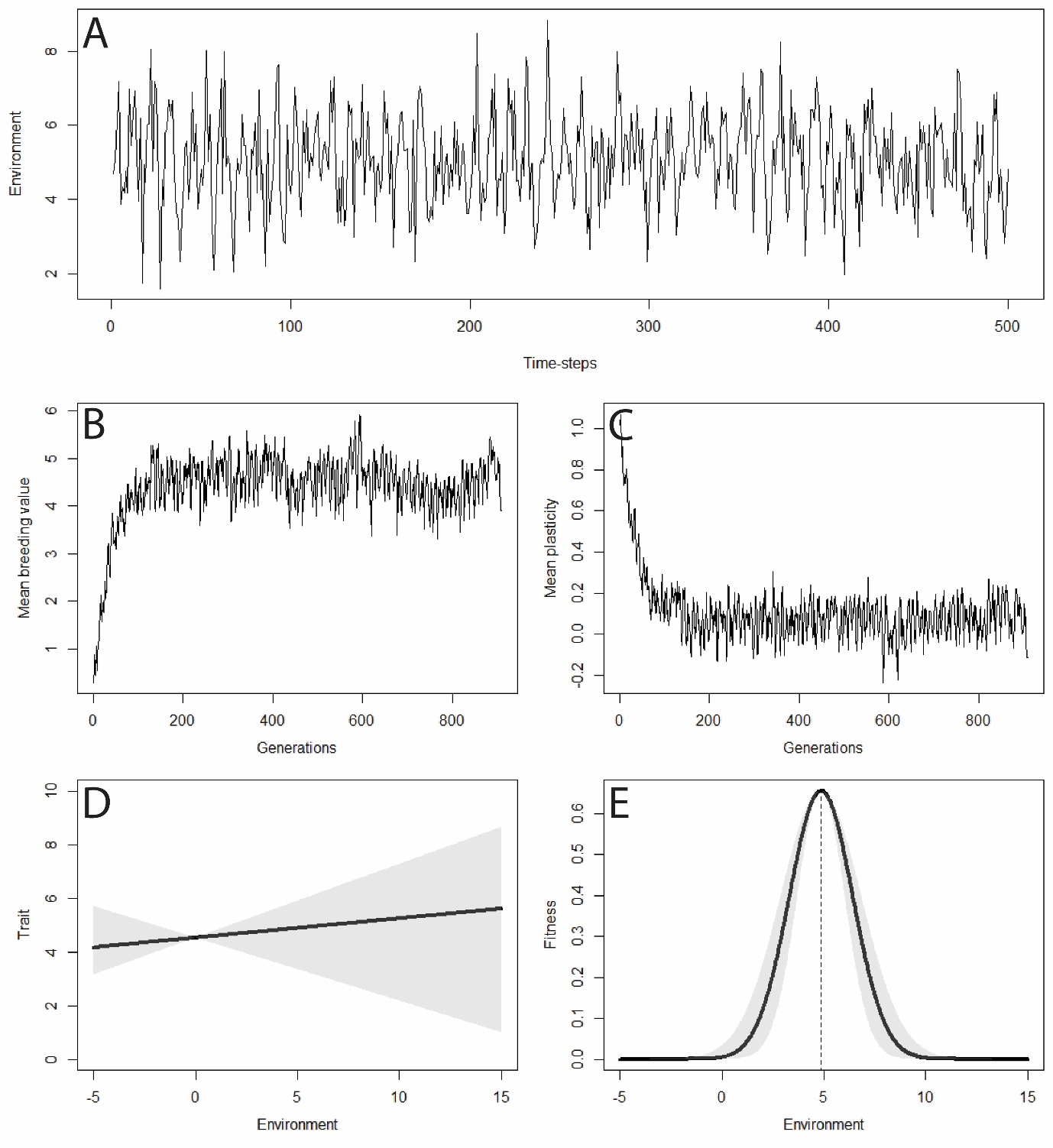

**Figure S1.** Evolution under the baseline environmental scenario (see above). (A) Excerpt of the environmental time series for which evolution was simulated. (B) Evolution of mean breeding value. (C) Evolution of mean plasticity. (D) Evolved trait reaction norm at equilibrium, with shaded area showing variation in mean plasticity around the equilibrium value. (E) Evolved tolerance curve at equilibrium, with shaded area showing variation in tolerance breadth around the equilibrium value. The dotted line is equivalent to $\varepsilon_{opt}$ in Figure 1.

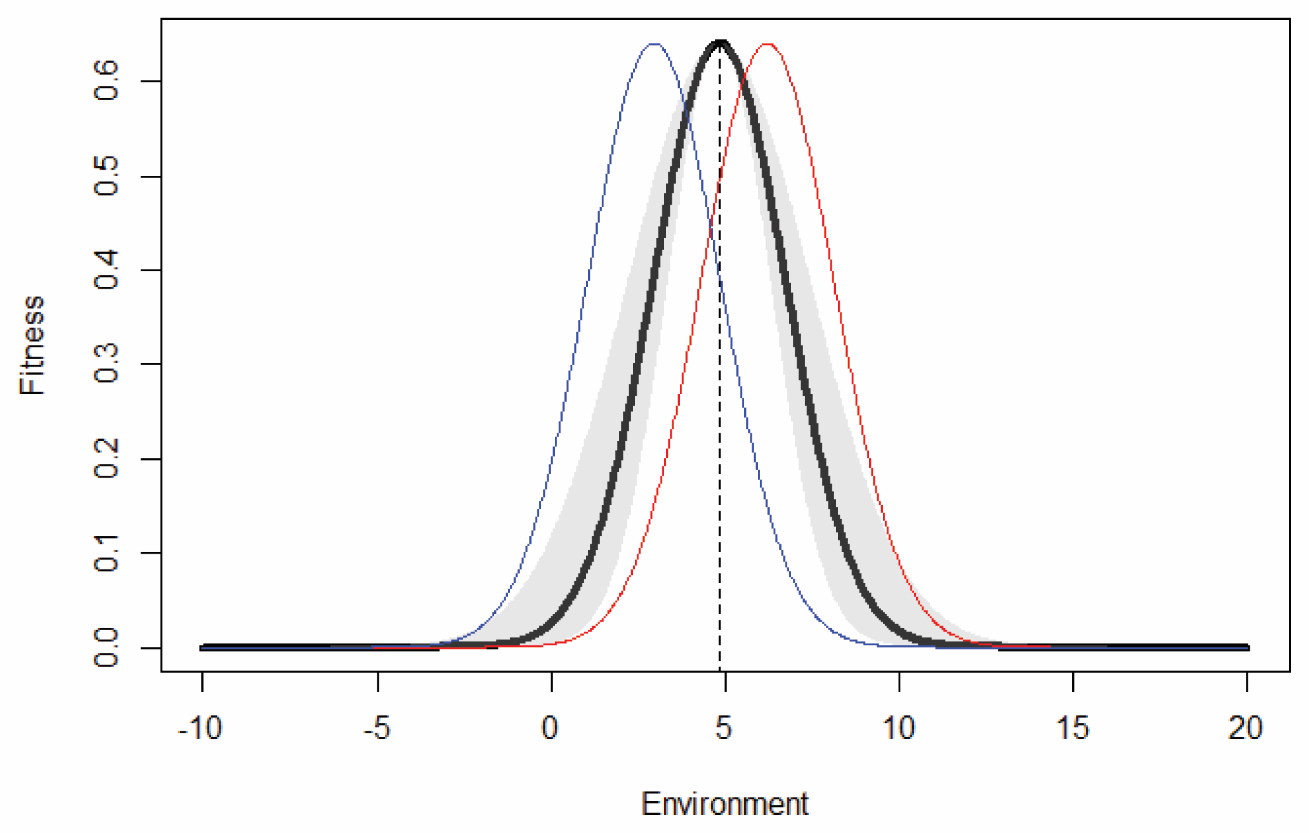

**Figure S2.** Variation in tolerance curve position (determined by the mean breeding value) and breadth (determined by mean plasticity) for the baseline environmental scenario. The black curve is the mean tolerance curve at dynamic equilibrium (generation 400 on), with shaded area showing variation in tolerance breadth predicted from maximum and minimum plasticities at equilibrium. The dotted line is equivalent to $\varepsilon_{opt}$ in Figure 1. Red and blue curves are predicted from maximum and minimum mean breeding values at equilibrium, respectively.

**Table S3.** Mean breeding values ($\bar{x}_{eq}$) and plasticities ($\bar{b}_{eq}$), with minima and maxima beneath, under changes in environmental mean, trend, cyclic change, noise, and noise colour (autocorrelation of noise). Diagonal elements are values after environmental components changed in isolation and off-diagonal elements are values after combined changes. Values were calculated at dynamic equilibrium (generation 400 onwards) and used to predict tolerance curves in Figure 4.

|  | **Change in mean** | **Trend** | **Cyclic change** | **Noise** | **Noise colour** |
| --- | --- | --- | --- | --- | --- |
| **Change in mean** | $\bar{x}_{eq}$ = 9.537  [9.045, 10.119]  $\bar{b}_{eq}$ = 0.035  [-0.192, 0.181] |  |  |  |  |
| **Trend** | $\bar{x}_{eq}$ = 16.035  [15.538, 16.502]  $\bar{b}_{eq}$ = 0.174  [0.081, 0.248] | $\bar{x}_{eq}$ = 12.299  [11.737, 12.834]  $\bar{b}_{eq}$ = 0.146  [0.024, 0.241] |  |  |  |
| **Cyclic change** | $\bar{x}_{eq}$ = 6.116  [5.101, 7.128]  $\bar{b}_{eq}$ = 0.340  [-0.029, 0.571] | $\bar{x}_{eq}$ = 9.379  [8.582, 10.077]  $\bar{b}_{eq}$ = 0.327  [0.113, 0.493] | $\bar{x}_{eq}$ = 1.909  [-0.112, 4.132]  $\bar{b}_{eq}$ = 0.545  [0.125, 0.824] |  |  |
| **Noise** | $\bar{x}_{eq}$ = 9.512  [8.404, 10.971]  $\bar{b}_{eq}$ = 0.030  [-0.043, 0.296] | $\bar{x}_{eq}$ = 12.276  [11.525, 12.921]  $\bar{b}_{eq}$ = 0.151  [-0.066, 0.311] | $\bar{x}_{eq}$ = 2.305  [-1.433, 6.137]  $\bar{b}_{eq}$ = 0.461  [-0.220, 0.912] | $\bar{x}_{eq}$ = 4.497  [1.924, 7.905]  $\bar{b}_{eq}$ = 0.064  [-0.555, 0.440] |  |
| **Noise colour** | $\bar{x}_{eq}$ = 9.772  [9.239, 10.287]  $\bar{b}_{eq}$ = 0.009  [-0.186, 0.189] | $\bar{x}_{eq}$ = 12.598  [12.018, 13.122]  $\bar{b}_{eq}$ = 0.124  [0.017, 0.244] | $\bar{x}_{eq}$ = 2.057  [-0.197, 3.560]  $\bar{b}_{eq}$ = 0.511  [0.184, 0.809] | $\bar{x}_{eq}$ = 5.219  [2.250, 7.869]  $\bar{b}_{eq}$ = -0.100  [-0.501, 0.279] | $\bar{x}_{eq}$ = 4.805  [3.434, 6.054]  $\bar{b}_{eq}$ = 0.010  [-0.235, 0.227] |

Baseline values for comparison: $\bar{x}_{eq}$ = 4.522 [3.318, 5.937], $\bar{b}_{eq}$ = 0.072 [-0.234, 0.274].

**Table S4.** Mean breeding values ($\bar{x}_{eq}$) and plasticities ($\bar{b}_{eq}$), with minima and maxima beneath, for locations in the case study. Values were calculated at dynamic equilibrium (generation 1,500 onwards) and used to predict tolerance curves in Figure 7.

| **Location** | **Mean breeding value (**$\bar{\boldsymbol{x}}$**)** | **Mean plasticity (**$\bar{\boldsymbol{b}}$**)** |
| --- | --- | --- |
| **Southern New South Wales** | $\bar{x}_{eq}$ = 19.697  [19.439, 19.922] | $\bar{b}_{eq}$ = 0.028  [-0.175, 0.194] |
| **Port Phillip Bay in Victoria** | $\bar{x}_{eq}$ = 14.092  [13.484, 14.448] | $\bar{b}_{eq}$ = 0.098  [-0.257, 0.377] |
| **Southwestern Tasmania** | $\bar{x}_{eq}$ = 13.976  [13.548, 14.238] | $\bar{b}_{eq}$ = 0.013  [-0.229, 0.195] |
